# Stress-induced adaptations in nucleus accumbens dopamine D1 receptor-expressing cells correspond to social avoidance behavior in male mice

**DOI:** 10.64898/2026.04.23.720476

**Authors:** Dominika Burek, William A. Carlezon

## Abstract

Stress can cause or exacerbate psychiatric illness, and effects on the transcription factor CREB within the nucleus accumbens (NAc) are critically involved. In rodents, stress-induced activation of NAc CREB produces elevations in dynorphin (DYN), an endogenous opioid expressed in dopamine D1-receptor (D1R)-expressing medium spiny neurons (MSNs). In turn, elevated DYN signaling produces features of mood and anxiety disorders via actions at kappa-opioid receptors (KORs). Although individual differences in stress sensitivity have been described—with some appearing susceptible and others resilient—the contribution of NAc DYN to these phenotypes is unclear. Here we examined relationships between social behavior and DYN in D1R-expressing MSNs in mice exposed to chronic social defeat stress (CSDS). We used quantitative (q)RNAscope to assess co-expression of genes encoding CREB (*Creb1*), D1Rs (*Drd1*), and DYN (*Pdyn*) within the NAc. To leverage individual variability, we performed regression analyses across all mice, revealing negative correlations between social interaction behavior and expression of *Drd1* and *Pdyn*, linking higher social avoidance with higher expression of these genes. There was no correlation with *Creb1*, suggesting stress-induced elevations in *Pdyn* depend on CREB activation (phosphorylation). These findings suggest that stress-induced elevations in D1R-associated DYN signaling within the NAc is a biomarker of susceptibility.

## INTRODUCTION

Stress is implicated in the etiology of mood and anxiety disorders including major depressive disorder (MDD) and post-traumatic stress disorder (PTSD) (Nestler and Carlezon, 2006; Ressler et al., 2022). These conditions share many core features, including diminished ability to experience pleasure or positive emotions (anhedonia), social avoidance, and sleep disruption (APA, 2013), suggesting a shared pathophysiology. The overlap in symptomatology illustrates potential benefits of using Research Domain Criteria (RDoC) principles (Cuthbert and Insel, 2013) to focus research on translationally-relevant features (e.g., social avoidance) that cut across multiple conditions rather than on narrow diagnostic categories that even highly skilled clinicians may struggle to differentiate in humans (Gros et al., 2012; Ressler et al., 2022). These symptoms are persistent, debilitating, and resistant to treatment, contributing to profound economic and social burdens (Greenberg et al., 1999; Birnbaum et al., 2010).

The biological basis of stress-related psychiatric disorders is unknown, which impedes the development of improved therapeutics. Stress can produce neuroadaptive responses (alterations in gene and protein expression) that contribute to persistent and often maladaptive changes in behavior (Nestler and Carlezon, 2006; Ressler et al., 2022). While these neuroadaptations occur throughout the brain, work focusing on the nucleus accumbens (NAc) provides strong evidence that stress produces changes in behavior via effects on CREB (cAMP response element binding protein) (Pliakas et al, 2001; Muschamp et al, 2011; Muschamp and Carlezon, 2013; Carlezon and Krystal, 2016), a transcription factor that regulates expression of myriad genes (Carlezon et al., 2005). A variety of stressors (forced swimming, footshock, social defeat) activate CREB in the NAc, which elevates expression of the opioid peptide dynorphin (DYN), which in turn causes depressive behaviors via activation of kappa-opioid receptors (KORs) (Muschamp and Carlezon, 2013; Carlezon and Krystal, 2016). A key mechanism of this effect is that increased DYN acts upon KORs expressed on the terminals and cell bodies of ventral tegmental area (VTA) dopamine (DA) neurons (Svingos et al, 1999; Svingos et al., 2001), producing inhibition of mesolimbic system function. The ability of KOR antagonists to relieve this inhibitory tone is a putative neural mechanism of their antidepressant- and anxiolytic-like effects (Carroll and Carlezon, 2013; Carlezon and Krystal, 2016). Increased CREB and KOR function are implicated in despair/learned helplessness (Pliakas et al., 2001), anxiety/avoidance behavior (Knoll et al., 2007), disrupted attention (Nemeth et al, 2010), anhedonia and fear (Muschamp et al., 2011), and sleep dysregulation (Wells et al., 2017; McCullough et al., 2021). Collectively, these studies suggest that a common pathophysiology within the NAc explains a broad assortment of features that are frequently co-morbid in stress-related disorders. An improved understanding of mechanism may enable development of new treatments that, by targeting the NAc and its specific cell populations, could dramatically improve outcomes by relieving numerous signs of stress-related illnesses simultaneously.

The NAc is the ventral aspect of the striatum, which is composed largely of MSNs that segregate into two major populations, defined by expression of DA D1 or D2 receptors (D1Rs and D2Rs, respectively) (Gerfen et al, 1990). Considerable evidence indicates that DYN in the NAc arises primarily from D1R-expressing MSNs. Specifically, seminal *in situ* hybridization studies showed that prodynorphin (*Pdyn*)—the gene encoding DYN—segregates with D1R– expressing MSNs, whereas proenkephalin (*Penk*)—encoding enkephalin—marks D2R-expressing MSNs. This pattern was subsequently confirmed in the human NAc (Hurd and Herkenham, 1995). More recent cell type–specific approaches in mice, including D1/D2 neuronal profiling (Lobo et al., 2006), translational profiling (TRAP) (Heiman et al., 2008), and single-cell transcriptomics (Saunders et al., 2018) consistently show that expression of *Pdyn* occurs almost exclusively in D1R-MSNs, with little-to-no expression in D2R-MSNs.

The present studies were designed to explore relationships between CREB and DYN expression within the NAc and stress-induced social avoidance behavior, a core feature of mood and anxiety disorders that can be studied in species ranging from mice to humans. Given convergent lines of evidence demonstrating that *Pdyn* in the NAc is predominantly restricted to D1R-expressing MSNs, we focused on the effects of stress on gene expression in this cell population. Using multiplex quantitative (q)RNAscope to enable simultaneous assessment of numerous transcripts (McCullough et al., 2018), we assessed expression and co-expression of *Creb1* (encoding CREB), *Drd1* (encoding D1Rs), and *Pdyn* in mice subjected to a 10-day chronic social defeat stress (CSDS) regimen and subsequently tested in open field social interaction (OFSI) tests (Donahue et al., 2014; Hisey et al., 2023). We performed this assessment in two ways: using a traditional approach where CSDS-exposed mice were segregated into stress-susceptible and resilient groups on the basis of social interaction behavior (Krishnan et al., 2007; Wells et al., 2017; McCullough et al., 2021), and by disaggregating the groups to explore relationships between these markers and behavior in individual mice, leveraging variability in each endpoint. We discovered significant group differences in *Drd1* and *Pdyn* expression using both approaches, with the disaggregated data revealing at an individual subject level strong correlations between levels of gene expression and social avoidance, a shared feature of MDD and PTSD.

## METHODS

### Mice

Subjects were adult male wild-type C57BL/6J mice obtained from Jackson Laboratory (Bar Harbor ME). To generate aggressors, virgin male and ovariectomized female Swiss Webster (CFW) mice were obtained at 8 weeks of age from Charles River (Shrewsbury, MA, USA). Mice were maintained in a temperature- and humidity-controlled vivarium under a 12-hour light/dark cycle. Food and water were available *ad libitum*. All housing and treatment procedures were approved by the McLean Hospital Institutional Animal Care and Use Committee following guidelines set by the National Institutes of Health.

### Chronic social defeat stress (CSDS)

All mice habituated to the vivarium for one week after arrival. Male CFW mice were pair-housed with ovariectomized female CFW mice for at least one week before separation and aggressor screening. Male CFW mice with an attack latency of <30 sec for 2 consecutive days were used for social defeat. Male C57 mice were subjected to a 10-day CSDS regimen, as described previously (Donahue et al., 2014; Wells et al., 2017; McCullough et al., 2021). Each day, a C57 was placed in the home cage of an aggressive CFW male for defeat for 5 min or until 50 bites were delivered. Mice were then separated in the CFW male’s home cage by a perforated Plexiglas divider that allowed for visual and olfactory contact without further physical interaction. C57 males were placed in a new CFW male’s home cage for each subsequent defeat.

### Open field social interaction (OFSI)

To identify susceptibility versus resilience to the CSDS paradigm, C57 mice were tested for social interaction in a square (45.7 x 45.7 cm) arena. The day after the tenth and final defeat session, each C57 was placed in the arena containing an empty wire cup for 150 sec before removal. A nonaggressive CFW male was then placed inside the wire cup and the C57 placed back into the arena. Motion was tracked and analyzed using EthoVision (Stoelting). Social interaction ratio was calculated as the amount of time spent in the social interaction zone (2 cm around the cup) when the CFW was present divided by the time spent when the CFW was absent.

### Quantitative (q)RNAscope

C57 mice were sacrificed 24 hr after OFSI, and brains were flash-frozen in isopentane on dry ice. The NAc was sectioned at 20 µm and RNAscope Multiplex Fluorescent V2 Assay (ACDBio) was conducted according to manufacturer instructions and previous studies (McCullough et al., 2018). Probes for *Creb1* (Mm-CREB-C2, Cat No. 496401-C2), *Drd1* (Mm-Drd1-C3, Cat No. 461901-C3) and *Pdyn* (Mm-Pdyn, Cat No. 318771) were hybridized to target RNA, then conjugated to Opal dyes (Akoya Biosciences) for fluorescent staining (Opal 690 for *Creb1*, and Opal 480 for *Drd1*, and Opal 570 for *Pdyn*).

### Confocal microscopy imaging

Images of each individual hemisphere were acquired on a Leica SP8 confocal microscope using a 20X objective and a 3×3 tilescan at a 1.0 zoom factor. Images were z-stacks of 10 steps at 0.5 microns per step using identical settings for laser power, detector gain, and amplifier offset calibrated to the brightest section.

### HALO image analysis

Images were quantified using the HALO imaging analysis software (V3.5, Indica) according to the manufacturer instructions (Halo 3.6 FISH-IF, v2.2). Probe copy intensity, used for determining the number of copies within a cluster, was set to the average intensity for cells expressing a single probe copy. DAPI and probe copy intensity, contrast threshold, signal minimum intensity, spot size, and segmentation aggressiveness were determined according to signal from positive control and noise from negative control sections.

### Statistical analyses

Counts of cells expressing any or a combination of probes were normalized to total cells analyzed. Probe copies were normalized to number of cells expressing any amount of probe. A cell was considered probe-positive if it contained at least 1 copy of the probe. Data from each hemisphere per animal were averaged. Gene expression was assessed in mice classified as susceptible or resilient and compared to controls using one-way ANOVAs followed by Tukey’s multiple comparisons tests. Sub-groups were then disaggregated and relationships between social behavior and gene expression across individual mice were assessed using simple correlation (Pearson’s) analyses, followed by linear regression comparisons with F-tests to assess group (defeated versus control) differences in slopes.

## RESULTS

The day after completion of the 10-day CSDS regimen (Day 11), defeated (DEF) mice and controls (CTRLs) were assessed in OFSI tests (**Fig. 1A**). As in previous studies, an SI ratio score of 1 was used as an index to quantify stress susceptibility (SI≥1, stress-resilient [RES]; SI≤1.0, stress-susceptible [SUS]) (Krishnan et al., 2007). When segregated into these classifications, a one-way ANOVA revealed significant group differences (F(2,12)=15.40, p=0.0005) (**Fig. 1B)**. *Post hoc* tests revealed that while there were no differences between RES and CTRL mice, SI ratios were significantly lower in SUS mice when compared to both CTRL (p=0.0021) or RES mice (p=0.0007). These findings confirm that our CSDS regimen produces reductions in SI in a sub-population of mice (SUS) that reflect increased social avoidance behavior (Berton et al., 2006; Krishnan et al., 2007), a core feature of mood and anxiety disorders (APA, 2013), while other mice (RES) appear unaffected.

**Fig. 1:**
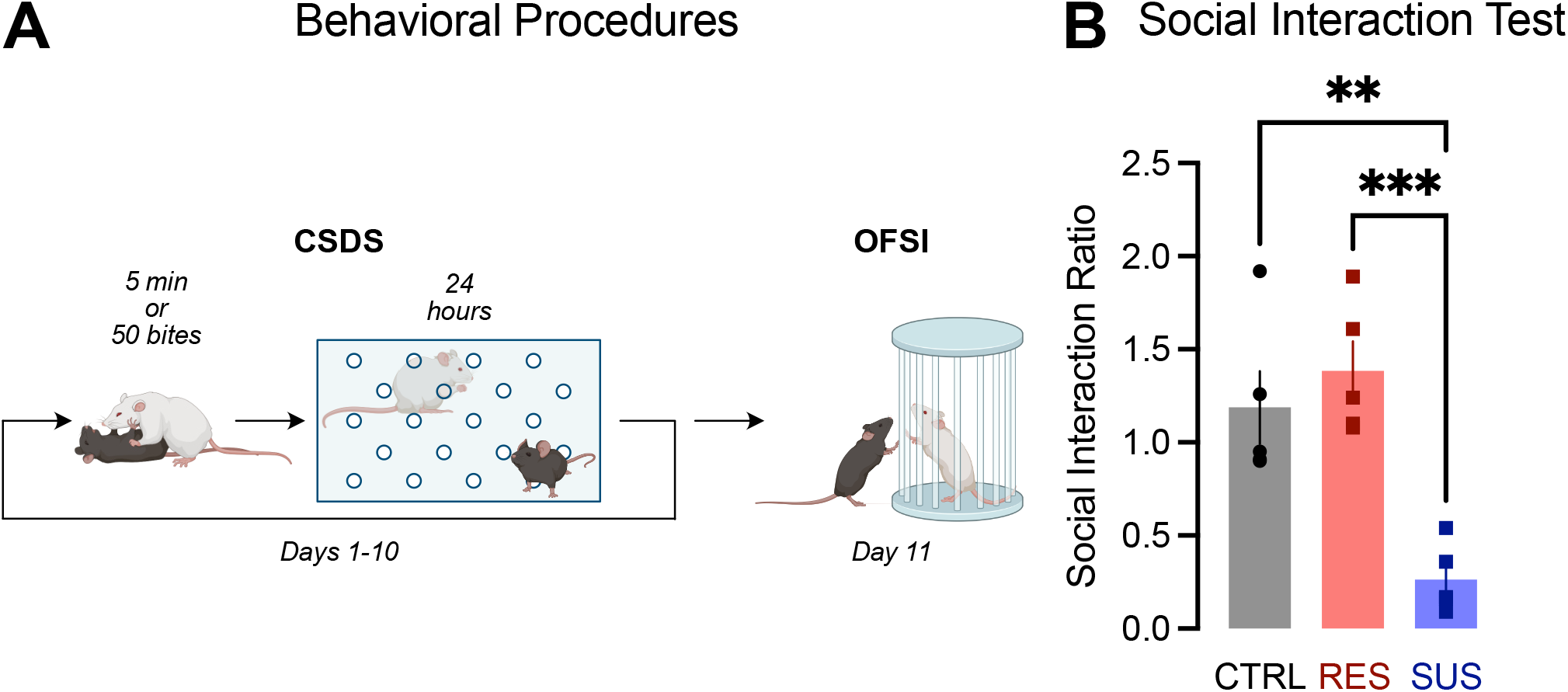
**(A)** Schematic of behavioral procedures involving chronic social defeat stress (CSDS) and open field social interaction (OFSI) tests. Adult male C57BL/6J mice were defeated by adult CFW male aggressors each day on 10 consecutive days. Each defeat session was followed by 24 hr of co-housing separated by a perforated plastic divider. On Day 11, social behavior was assessed in OFSI tests. **(B)** When defeated mice were segregated into subgroups on the basis of their OFSI behavior, those categorized as stress-susceptible (SUS) had social interaction ratios that were significantly lower than controls (CTRLs) and those categorized as stress-resilient (RES), indicating increased social avoidance. ^*\*\**^*P*<0.01, ^***^*P*<0.001, Tukey’s post hoc tests, N’s=5/group.

We first performed qRNAscope *in situ* hybridization on 20-µm sections of NAc from each mouse to assess levels of *Creb1, Drd1*, and *Pdyn*. Each hemisphere was imaged and analyzed separately, and data from each was averaged per mouse. Reliable expression of all probes was detectable (**Fig. 2A-B**). We then aggregated the DEF mice according to their phenotypic designations (RES, SUS), and assessed group differences in the number of cells expressing the transcripts. Overall, CSDS had no effect on any of these endpoints: there were no group differences in total the number of cells expressing DAPI (used to normalize quantification) (F[2,12]=2.531, p=0.1210, not significant [ns]), *Creb1* (F[2,12]=0.3508, p=0.7111, ns), *Drd1* (F[2,12]=1.760, p=0.2137, ns), *Pdyn* (F[2,12]=0.3592, p=0.7055; ns), or co-expressing *Drd1+Pdyn* (F[2,12]=0.6527, p=0.5381, ns) (data not shown). Findings were consistent with previous reports, including ubiquitous expression of CREB (∼99% of cells) (Carlezon et al., 2005), and virtually complete (∼95%) overlap in co-expression of *Drd1* and *Pdyn* (Gerfen et al., 1990). Together, these findings suggest that CSDS does not cause fundamental shifts in the number of MSNs co-expressing D1Rs and DYN.

**Fig. 2:**
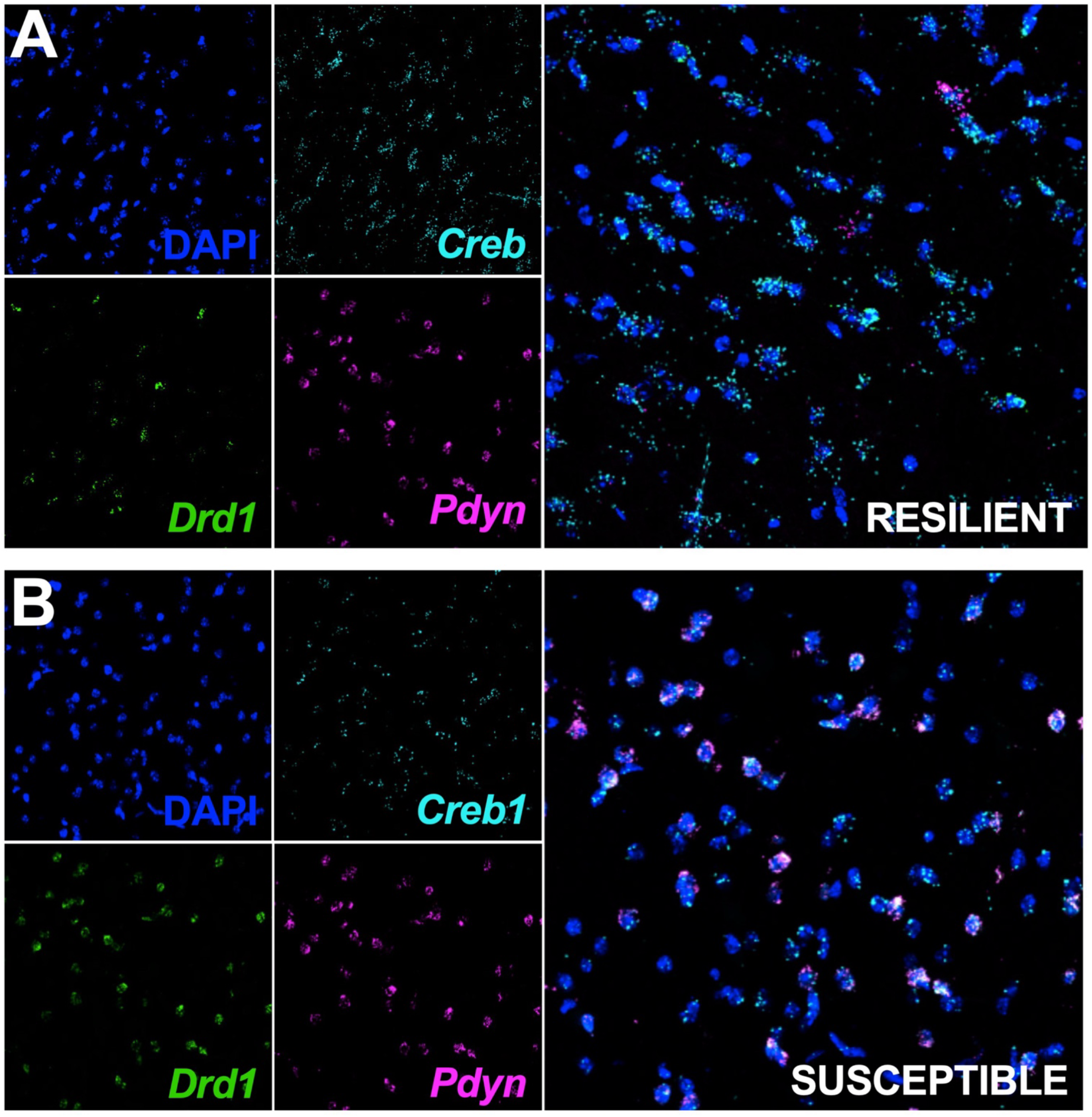
Multiplex quantitative (q)RNAscope to assess group differences in expression of *Creb1, Drd1*, and *Pdyn* the nucleus accumbens (NAc). DAPI stain labels all cells. **(A)** Images from a representative stress-resilient (RES) mouse depicting each of the individual channels (DAPI=blue, *Creb1*=cyan, *Drd1*=green, *Pdyn*=magenta) (left) and overlay (right). **(B)** Images from a representative stress-susceptible (SUS) mouse, which on average had higher expression of *DrD1* and *Pdyn* per labeled cell.

Next, we assessed group differences on the amount of transcript—reflected by the number of probe copies—per labeled (+) cell. There were no group differences in the amount of *Creb1* transcript per *Creb1+* cell (F[2,12]=0.2426, p=0.7883, ns; **Fig. 3A**), consistent with prior evidence indicating that dynamic regulation of CREB occurs via treatment-related changes in phosphorylation rather than at the level of gene and protein expression (Carlezon et al., 2005). However, there were group differences in the amount of *Drd1* per *Drd1+* cell (F[2,12]=9.718, p=0.0031 **Fig. 3B)**, with higher expression in SUS mice compared to CTRL (p=0.0138) and RES mice (p=0.0036). Likewise, there were group differences in the amount of *Pdyn* per *Pdyn*+ cell (F[2,12]=4.864, p=0.0284, **Fig. 3C**), with higher levels of expression in SUS compared to RES mice (p=0.0384). Together, these findings suggest that CSDS produces neuroadaptive responses within the NAc that increase both sensitivity to the excitatory (G_S_-coupled) effects of DA and corresponding DYN release in SUS but not RES mice.

**Fig. 3:**
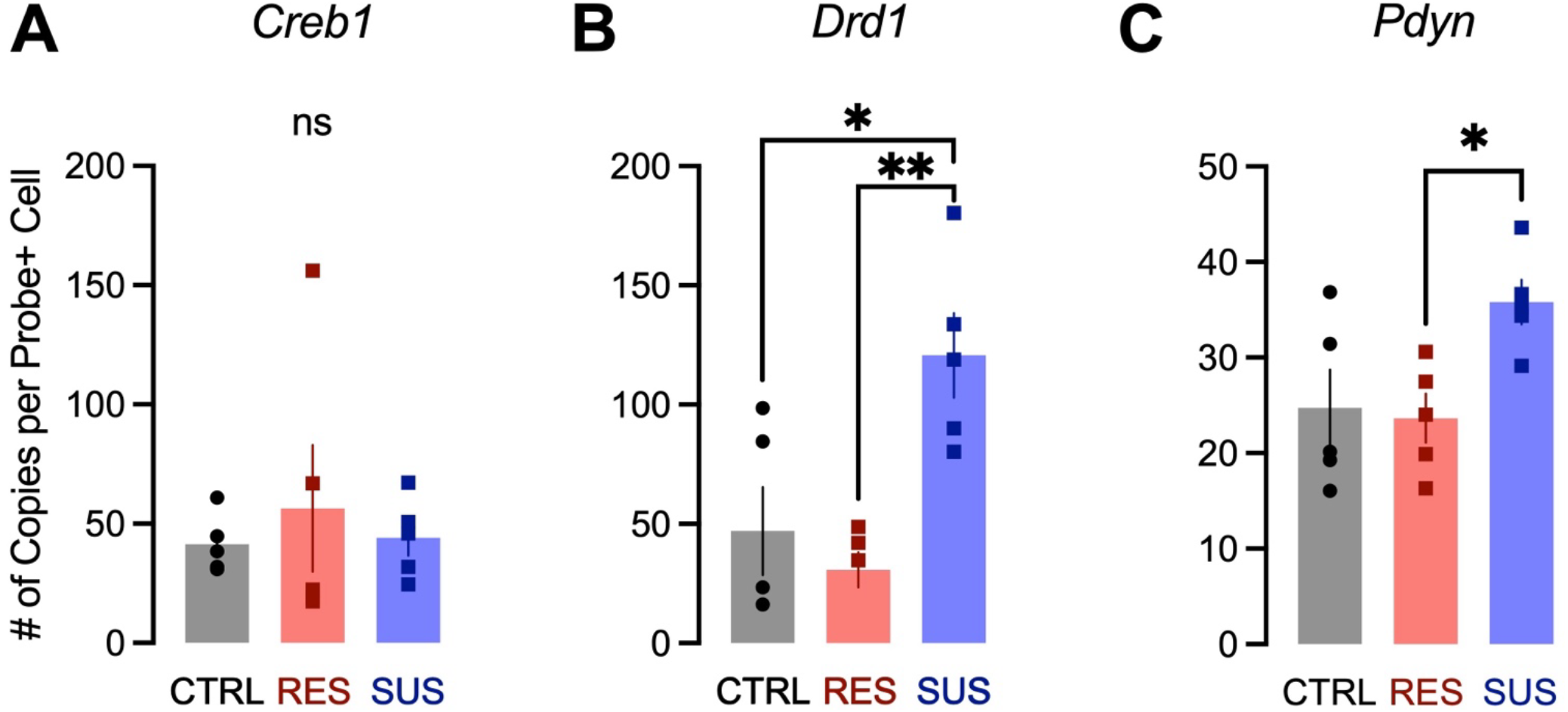
Quantification of *Creb1, Drd1*, and *Pdyn* levels in the NAc, as reflected by the number of probe copies per labeled cell. While there were no group differences in *Creb1* **(A)**, *Drd1* levels were higher in SUS mice than in RES mice or CTRL **(B)**. Similarly, *Pdyn* levels were higher in SUS mice than in RES mice, which did not differ from CTRLs **(C)**. ^*\**^*P*<0.05, ^**^*P*<0.01, Tukey’s post hoc tests.

Finally, we disaggregated the groups to leverage individual differences in social behavior and gene expression, enabling assessment of correlations across the entire continuum of endpoints. There were no correlations between social interaction and *Creb1* copies in *Creb1+* cells in DEF (SUS+RES combined) mice (Pearson r=-0.6210, p=0.2636, ns) or CTRLs (r=0.1801, p=0.6185, ns) (**Fig. 4A**), again suggesting that stress has little effect on CREB transcript levels. There was no significant difference in slopes between the groups (F[1,11]=0.3798, p=0.5503, ns). In contrast, there was a significant negative correlation between social behavior and *Drd1* copies per *Drd1*+ cells (r=-0.7524, p=0.0120), with lower social approach/higher avoidance in DEF mice correlating with higher *Drd1* expression (**Fig. 4B**). There was no correlation in CTRL mice (r=0.5753, p=0.3102, ns), nor was there a group difference in slopes (F[1,11]=0.04158, p=0.8421, ns). Similarly, there was a significant negative correlation between social behavior and *Pdyn* copies per *Pdyn*+ cell (r=-0.7822, p=0.0075), with lower social approach/higher avoidance in DEF mice correlating with higher *Pdyn* (**Fig. 4C**). There was no correlation in CTRL mice (r=0.6372, p=0.2475, ns), but there was a group differences in the slopes (F[1,11]=8.630, p=0.0135). Together, these findings link higher social avoidance with higher expression of these genes in DEF mice, regardless of phenotypic classification as SUS or RES.

**Fig. 4:**
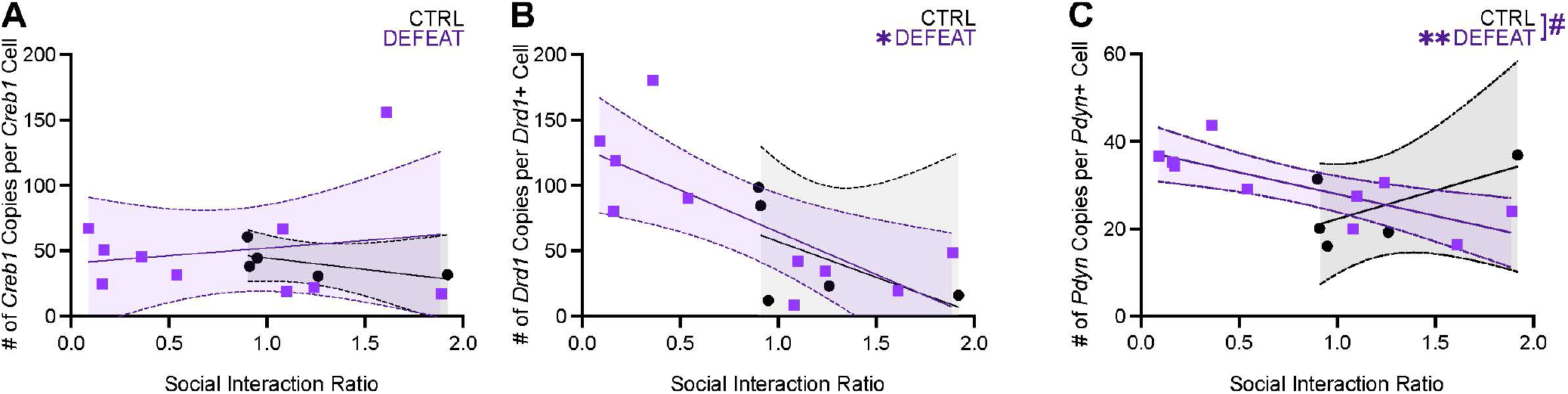
Assessment of correlations between SI behavior and gene expression when RES and SUS groups are consolidated into a single defeated (DEF) condition and compared to controls. **(A)** There was no correlation between SI behavior and *Creb1* levels in either DEF mice or CTRLs, nor were there differences in slope. **(B)** There was a significant negative correlation between *Drd1* levels and social behavior in DEF mice, with a higher of probe copies correlating with lower SI (i.e., higher social avoidance). The correlation was not significant in CTRLs, nor did the slopes differ. **(C)** Similarly, there was a significant negative correlation between *Pdyn* levels and social behavior in DEF mice, with a higher of probe copies correlating with lower SI. There was no correlation in CTRLs, and the slopes were significantly different between groups. ^*\**^*P*<0.05, ^**^*P*<0.01, Pearson’s tests; #*P*<0.05, F-tests.

## DISCUSSION

Here we examined relationships between social behavior and DYN in D1R-expressing MSNs in mice exposed to CSDS, using multiplex (q)RNAscope to assess expression of *Creb1, Drd1*, and *Pdyn* within the NAc. Defeated mice were first phenotyped on the basis of their social behavior, with those classified as SUS showing lower social interaction (and thus higher social avoidance) than controls, and those classified as RES showing social interaction similar to controls. In mice categorized in this way, we saw no group differences in the number of NAc cells expressing *Creb1, Drd1, Pdyn* or *Drd1+Pdyn*, suggesting that CSDS does not cause fundamental changes in the distribution of these transcripts. However, we identified group differences on the amount of transcript—reflected by the number of probe copies—per labeled cell. Specifically, SUS (socially-avoidant) mice had higher levels of *Drd1* in *Drd1+* cells and *Pdyn* in *Pdyn+* cells, suggesting that CSDS can produces increases in D1R and DYN tone and that occurrence of such changes contributes to this phenotype. To leverage individual variability, we performed regression analyses across all mice, combining SUS+RES into a DEF condition and comparing it to controls. These analyses revealed negative correlations between social interaction behavior and expression of *Drd1* and *Pdyn*, linking higher social avoidance with higher expression of these transcripts across a continuum of behavior and gene expression levels. No group differences or correlations were observed with *Creb1*, suggesting stress-induced elevations in *Pdyn*—a known CREB target gene (Carlezon et al., 1998; Carlezon et al., 2005)—may depend more on CREB activation (phosphorylation) than levels of the gene or protein. When considered together, these findings suggest that stress-induced elevations in D1R-associated DYN signaling within the NAc is a biomarker of susceptibility.

Our findings are consistent with a broad literature linking increased NAc DYN and depressive behavior. Seminal work showed that elevated CREB function in the NAc produces increases in DYN expression, which leads to a variety to depressive- and anxiety-like phenotypes (Carlezon et al., 1998; Pliakas et al., 2001; Muschamp and Carlezon, 2013; Carlezon and Krystal, 2016). These effects are reproduced, at least in part, by administration of KOR agonists (which mimic DYN actions at KORs) directly into the NAc (Bals-Kubick et al., 1993; Muschamp et al., 2011). Likewise, administration of KOR agonists to humans causes signs of mood and anxiety disorders (Pfeiffer et al., 1986; Walsh et al., 2001). The depressive effects are typically attributed to elevated DYN expression in D1R-MSNs, considering evidence that links this peptide and cell type in striatal regions including the NAc (Gerfen et al., 1990; Hurd and Herkenham, 1995; Lobo et al., 2006; Heiman et al., 2008; Saunders et al., 2018). A wide variety of stressors—including forced swimming, footshock, and drug withdrawal—activate CREB via phosphorylation, identifying a shared molecular cascade that in humans could create psychopathology while creating interest in the use of KOR antagonists to treat depression and anxiety (Muschamp and Carlezon, 2013; Carlezon and Krystal, 2016). Interestingly, there is evidence that increased activity of NAc D1-MSNs before stress predicts resiliency in mice (Muir et al., 2018). In the present studies, we found elevated *Drd1* expression in the NAc of SUS rather than RES mice. While these findings may initially seem inconsistent, we also found elevated *Pdyn* in SUS mice. When considered together, these studies raise the possibility that elevated activity of D1R-MSNs is generally associated with resilience, particularly when it precedes stress exposure that produces coincident elevations in DYN. Applying this conceptualization to the present data, CSDS-induced elevations in *Drd1* in SUS mice could represent a homeostatic neuroadaptation that is overshadowed by the depressive effects of elevated D1-MSN function-related increases in DYN release. As such, elevated *Pdyn* creates a mixed signal in which factors associated with resilience co-occur with factors associated with susceptibility: in stressed individuals, elevations in D1 signaling that were protective before stress exposure now also produce secondary effects (e.g., enhanced inhibition of mesolimbic DA neurons via elevated DYN tone) that promote depressive behaviors including social avoidance. This mixed effect may contribute to difficulties identifying biomarkers (e.g., elevated NAc D1R expression) that can predict susceptibility to stress, particularly when an individual’s stress history is not known. Our studies suggest that elevated NAc DYN represents a yet-to-be explored biomarker of stress susceptibility, since its presence or absence affects the consequences of increased D1R-MSN activity (Muir et al., 2018). This hypothesis could be tested directly in humans using brain imaging.

These present proof-of principle studies have some limitations, while identifying several areas for follow-up research. Foremost, it is important to perform complementary studies in females, particularly considering that depressive illness is twice as prevalent in women as in men (Paykel, 1991; Desai and Jann, 2000) and reports of sex differences in KOR system function in rodents (Russell et al., 2014; Conway et al., 2019). It is notoriously difficult to demonstrate CSDS-induced social avoidance in female mice, although recent studies suggest that this can be mitigated by procedural changes that increase aggression during the defeat sessions (Newman et al., 2019), or by assessing other behavioral endpoints (Schuler et al., 2025). Complementary studies in female are difficult to design without reliable and well-validated behavioral endpoints to use in combination with either the traditional SUS/RES dichotomy or disaggregated (individual) data. Additionally, the roles of other NAc cell types in DYN/KOR-related processes are currently unclear. For example, there is evidence that microglia express KORs and, under inflammatory conditions, DYN (Missig et al., 2022; Li et al., 2017). Presence of these markers in non-neuronal cells—where they may be regulated differently than in neurons—can make it difficult to devise circuit models, particularly without being able to identify the cell types contributing to the DYN/KOR signal. Future work is needed to more clearly characterize DYN expression in neuronal and non-neuronal cell populations under baseline conditions and in response to stress. Finally, it will be important to extend these types of studies to other translationally-relevant behavioral domains (e.g., anhedonia) that are also core features of stress-related illness and characterize some subtypes of depression (Dodd et al., 2021). In mice, there are currently few well-validated behavioral endpoints that are conducive to segregation into the RES/SUS dichotomy, although use of disaggregated data across numerous endpoints may offer the ability to leverage individual variability to sharpen predictions and enable development of translationally-relevant biomarkers for psychiatric illness. Regardless, the present studies strengthen the link between DYN signaling in the NAc and stress-induced development and expression of depressive behaviors.

## FUNDING

Supported by R01MH063266 (to WAC).

## DISCLOSURES

Within the past 3 years, WAC has served as a consultant for AbbVie, Neumora, and Psy Therapeutics, and received sponsored research agreements from AbbVie and Delix. None of the other authors report disclosures.

